# Rheumatoid arthritis patients express a skewed repertoire of polyclonal, hypomutated B-cell receptors

**DOI:** 10.1101/771949

**Authors:** Graeme J.M. Cowan, Katherine Miles, Lorenzo Capitani, Sophie S.B. Giguere, Hanna Johnsson, Carl Goodyear, Iain B. McInnes, Scottish Early Rheumatoid Arthritis Inception cohort Investigators, Steffen Breusch, David Gray, Mohini Gray

## Abstract

**Objectives:** The success of B cell depletion therapy in rheumatoid arthritis (RA) therapy testifies to their importance in disease pathogenesis, but the precise B cells mediating this are unclear. For example, it is unknown if RA patients predominantly express a limited number of circulating clonally expanded populations of B cells with highly mutated B cell antigen receptors (BCRs) that would constitute a shared antigen driven response.

**Methods:** To address this, we have undertaken the largest study to date utilising next generation sequencing (NGS), to identify the full length of the peripheral blood BCR sequences from the antigen-binding heavy chain. Between 25,000 to 200,000 BCR sequences per patient were analysed from 127 newly diagnosed RA patients, 16 heathy controls, 16 RA patients with established disease and 8 paired blood and synovial samples. This was complemented with B cell subset analysis from an additional 64 RA patients and 22 healthy controls.

**Results:** RA patients expressed a significantly higher percentage of circulating poorly mutated polyclonal IgG^+ve^ variable heavy (IgG-Vh) BCR sequences, both at the time of diagnosis and following treatment. These sequences resided predominantly within TNF-alpha secreting IgG^+ve^CD27^−ve^ B cells, that were expanded in RA peripheral blood and enriched in the rheumatoid synovium. Surprisingly, peripheral and synovial B cell repertoires of RA patients are quite distinct, sharing very few IgG sequences.

**Conclusions:** This is the first report to conclusively establish that a substantial component of the peripheral B cell repertoire in RA consists of polyclonal hypomutated IgG^+ve^ BCRs that may play a critical role in driving an autoimmune mediated inflammation.

## Introduction

Rheumatoid arthritis (RA) is the commonest autoimmune inflammatory arthritis, affecting up to 1% of the world’s population (1, 2). It is characterised by chronic systemic inflammation that targets the synovial joints leading to progressive joint damage and disability. Whilst pathogenesis is incompletely understood, the pivotal role of B cells is supported by the efficacy of B cell depletion therapy (BCDT) in the majority of treated patients (2–4). Auto-antibodies binding to post-translationally modified epitopes [e.g. citrullinated protein antigens (ACPA)] or the constant region of other immunoglobulins [called rheumatoid factor (RF)] (5), augment the generation of immune complexes. However good clinical responses to BCDT do not always correlate with a reduction in the titre of ACPA(6, 7), implying that ACPA may originate from long lived plasma cells, that are not targeted by anti-CD20 antibody therapy (8). The promotion of inflammation by self reactive B cells is likely to extend well beyond these specifcities. Understanding this would lead to more specific RA therapies and the hope of an eventual cure.

The advent of accessible, high throughput sequencing technologies has enabled formal evaluation of differences between the B cell repertoire of healthy and diseased individuals. To date, studies characterising the B cell repertoire of RA patients have been limited in number, sample size or sequencing depth (9). An understanding of the expressed BCR repertoires in RA patients would provide a more complete understanding of disease pathogenesis. We hypothesized that RA patients would have expansions of circulating pathogenic B cells at diagnosis, that could also be detected as a conserved BCR signature in established disease. Utilising next generation sequencing (NGS) of peripheral blood and synovial B cells, we sequenced the repertoire of expressed BCRs, focusing on the main antigen binding IgG variable heavy (IgG-Vh) region. We assessed 127 newly diagnosed RA patients, 16 patients with established RA and 8 paired blood and synovial samples. In addition we phenotyped B cells from an additional 64 RA patients and 22 healthy controls. Surprisingly, RA patients expressed significantly more IgG^+ve^ BCR sequences with fewer than five mutations (which we refer to hereafter as hypomutated or IgG^hypoM^). The hypomutated Vh-IgG BCRs were polyclonal and originated from TNF-alpha secreting IgG^+ve^CD27^−ve^ cells, which were significantly increased in the circulation. Unexpectedly, we detected a minimal sharing of sequences either between patients or between the synovium and peripheral blood of the same patient.

## Materials and Methods

### Ethical review and donor selection criteria

The use of human samples for cohorts 1, 3 and 4 was approved by the South East Scotland Bioresource NHS Ethical Review Board (Ref 15/ES/0094). Ethical permission to collect samples donated to the SERA inception (cohort 2) was approved by the West of Scotland Local Research Ethics Committee (Ref 10/S0703/4) as previously described(10). Informed consent was obtained from all study participants prior to sample collection. Patient cohorts are described in supplementary Tables 1-4.

### Patient and Public Involvement

Patients were not involved in the design of this study. Patients will be informed of the results of the study through the activities of the Centre for Inflammation Research public engagement events (https://www.edweb.ed.ac.uk/inflammation-research/information-public) and patient groups, including NRAS.

### Flow Cytometry & FACS sorting

Lymphocytes were stained in PBS with 1% FCS for 20 minutes at 4 ^o^C. A BD Aria II was used for flow sorting and a BD LSRII was used to collect data. All analysis was performed using FlowJo software. Debris and dead cells were excluded using FSC-SSC. Doublets were excluded using both FSC and SSC singlet gating. A list of antibody reagents is shown in supplementary table 5. Intracellular staining was performed, following stimulation (for 4.5h) with PMA [Sigma (20 ng/ml)] & Ionomycin [Sigma (1µg/ml)]. After 1h of stimulation Brefeldin A [Sigma (1µg/ml)] was added for the remaining 3.5 hrs. Surface staining was performed before fixation and permeabilisation using a Cytofix/Cytoperm kit (BD Biosciences).

### Cell purification

Peripheral blood mononuclear cells (PBMC) were prepared from citrated blood samples using Ficoll^®^ Paque Plus density centrifugation (GE Healthcare). Synovial tissue was digested for 2h at 37 ^0^C in 1 mg ml^−1^ Collagenase 1 (Sigma-Aldrich). PBMC and synovial tissue were enriched for B-cells using either anti-CD19 or anti-CD20 magnetic beads (Miltenyi Biotech) (see Supplementary table 6).

### B cell repertoire sequencing

Supplementary table 4 specifies the individual amplification strategies employed for samples. mRNA was purified using mRNA direct kit (Life Technologies), or total RNA was purified using either a Direct-zol total RNA kit (Zymo Research) or a Paxgene blood RNA kit (Qiagen). The amplification protocol, immune repertoire analysis and B cell clone lineage tree construction is provided in the online supplementary methods.

### Statistical analysis and data visualisation

Before performing inferential statistical tests, data were assessed for conformity to the assumptions of the test used. The assumption of normality of data was visually assessed using the Q-Q plot method, generated using the StatsModels Python package. Prism 6.0 (GraphPad Software Inc.) or the scipy.stats Python package (11) was used to perform all Student’s T tests, Mann-Whitney and Kruskal-Wallis non-parametric tests and Dunn’s post hoc test was run using the scikit-posthocs module (12). For analyses involving multiple pairwise comparisons, p-value adjustment was performed using the Holm-Šídák method (13). Two-tailed p-values are given in all cases. All plots were drawn with Prism (Graphpad Software Inc.) or with the Matplotlib (14) or Seaborn Python packages, and data processing used the Pandas package (15). Use of the ± following mean values indicates the 95% confidence interval.

## Results

### RA patients express a higher frequency of hypomutated IgG B cell receptors within peripheral blood

200,000 highly purified CD19^+ve^ B cells isolated from the peripheral blood of each of 14 newly diagnosed treatment-naïve seropositive early rheumatoid arthritis (ERA) patients (cohort 1/Supplementary Table 1) and 16 healthy controls were sequenced by NGS. The degree of mutation of the IgG heavy chain (IgG-Vh) was calculated by assessing the number of nucleotide mismatches between each sequence read and the closest predicted germline V segment sequence. Whereas the number of mutations per IgG read approximated to a symmetrical distribution in healthy individuals, the distribution of mutations in RA donors was skewed by the presence of an increased frequency of poorly mutated IgG sequences [Figure 1A-B/ Supplementary figure 2]. This observation was confirmed in a larger cohort of 113 newly diagnosed, DMARD naïve, seropositive ERA patients, selected from the Scottish Early Rheumatoid Arthritis (SERA) inception cohort(16) (cohort 2/Supplementary Table 2). A bimodal distribution of IgG-Vh mutation counts was observed in the IgG-Vh sequences with the first peak representing poorly mutated IgG sequences [Figure 1C]. To establish if this population persisted in patients with established RA (ESRA), we sequenced the IgG-Vh sequences from 16 ESRA donors (cohort 3/Supplementary Table 3). Analysis of the mean IgG mutation count per read showed that there were fewer IgG-Vh mutations in both of the ERA and ESRA cohorts, compared to healthy control donors [Figure 1D]. A further 12 paired samples taken 6 months following DMARD therapy (Supplementary Table 2) showed that the mean number of IgG-Vh mutations was only slightly increased [16.3 ± 2.4 compared with 12.8 ± 1.9 at diagnosis] (p< 0.006, Wilcoxon signed-rank test). The skewed distribution of IgG-Vh mutation counts in ERA donors were the result of an increased frequency of IgG sequences with fewer than 5 mutations. Indeed, the mean percentage of the IgG repertoire composed of fewer than 5 V-segment mutations (hypomutated IgG sequences [IgG^hypoM^]) in all three RA cohorts was significantly higher than healthy controls (mean for ERA cohorts 1 and 2 was 11.8% and 15.9% respectively, and for ESRA 18.5% compared to the mean for healthy controls of just 3.4%) [Figure 1E]. We did not detect a correlation between the frequency of IgG^hypoM^ and the degree of joint inflammation at the time of RA diagnosis as measured by the disease activity 28 joint score (DAS28) [Supplementary Figure 2B], which makes it unlikely that the presence of IgG^hypoM^ was simply a result of more marked inflammation.

**Figure 1.**
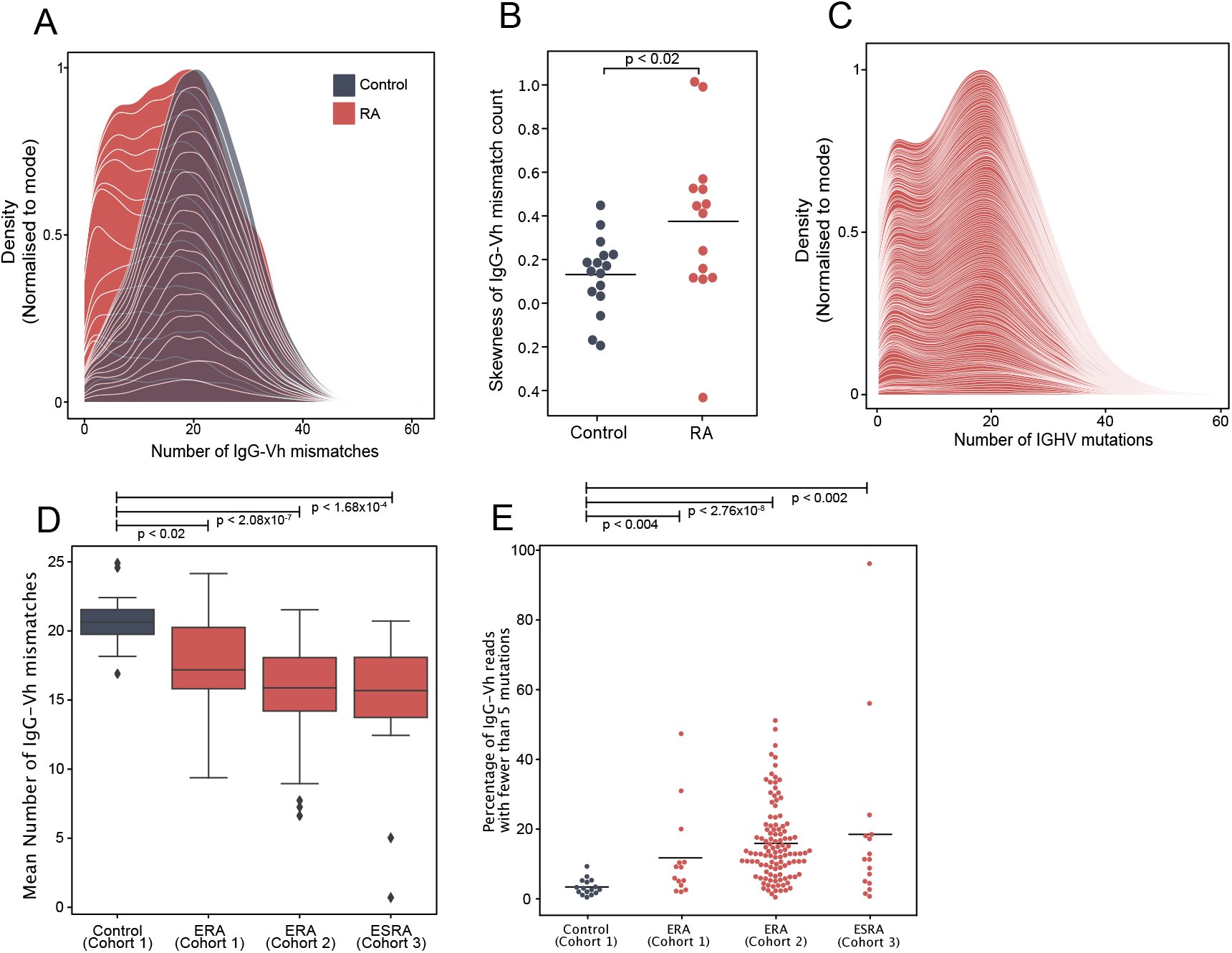
A. Distribution of the number of IgG-Vh mismatches per sequencing read for DMARD naïve early RA patients (ERA) (n=14) and healthy control donors (n=16). Individual density plots are stacked to indicate the overall distribution across all samples in each group. Maximum cumulative density values for each group are normalised to the mode to facilitate inter-group comparison. B. Skewness of IgG mutation distributions from RA patients (n=14) and healthy control groups (n=16). Horizontal lines denote the arithmetic mean skewness for each group. P value shown was calculated using Mann-Whitney U test. C. Distribution of the number of V segment mismatches per sequencing read for ERA patients [cohort 2, n=113]. Individual density plots are stacked to indicate the overall distribution across all samples in each group. D. Mean IgG-Vh mismatches for control donors (n=16), ERA donors from cohorts 1 and 2 (n=14 and n=113 respectively) and ESRA from cohort 3 (n=16). P values are generated by Kruskal-Wallis test with Dunn’s post-test to compare the means for each RA group with the control donor group. E. Percentage of IgG reads with fewer than 5 mutations for healthy donors from cohort 1 (n=16), ERA donors from cohort 1 and cohort 2 (n=14 and n=113) and ESRA donors from cohort 3 (n=16).

### Hypomutated sequences are distributed throughout the IgG repertoire

A potential explanation for the increased frequency of IgG^hypoM^ in RA donors could be mono- or oligo-clonal expansion of IgG B cell clonotypes with few mutations. However, this was unlikely given that the IgG^hypoM^ sequences were not restricted to particular IGHV allele families [Figure 2A and supplementary figure 3]. The IGHV4-34 gene segment, that conveys self-reactivity and is strongly associated with a failure of B cell tolerance and autoimmunity(17–19) is censored at multiple check points in healthy individuals(20). Yet, this was significantly higher in the IgG^+ve^ B cells from 113 DMARD naïve RA patients (2.41%, 95% C.I. 1.98-2.83%) compared to healthy controls 0.65%, (95% C.I. 0.50-0.79%) [Figure 2B]. Somatic mutations within the IGHV4-34 gene reduce self-reactivity(21, 22) but the IGHV4-34 allele in RA IgG^+ve^ BCRs expressed significantly fewer mutations than the healthy controls (p<7.31×10^−7^) [Figure 2C]. This demonstrates that RA patients express considerably more BCRs that utilize a poorly mutated IGHV4-34 allele. The IGHV4-34 allele is unusual in that it contains an Ala-Val-Tyr (AVY) motif (within the framework 1 region) responsible for the self-reactivity towards I/i carbohydrate antigens (18, 23, 24). There was no detectable difference in the mutation rate of AVY motifs between RA or healthy control donors in sequences of either the IgG or IgM isotype [Figure 2D]. The Asn-X-Ser N-glycosylation site (NHS) in the CDR2 region is associated with binding to commensal bacteria by innate like B cells(25) and is usually mutated in IgG^+ve^ B cells(22). In RA patients the proportion of IGHV4-34 IgG sequences where the NHS N-glycosylation motif was still intact was significantly higher (46.8% [IQR:33.6%-59.9%]) compared to healthy control donors (26.9% [IQR:21.1%-32.6%]) [Figure 2E]. Hence RA patients express significantly more polyclonal, hypomutated self-reactive IgG^+ve^ BCRs.

**Figure 2.**
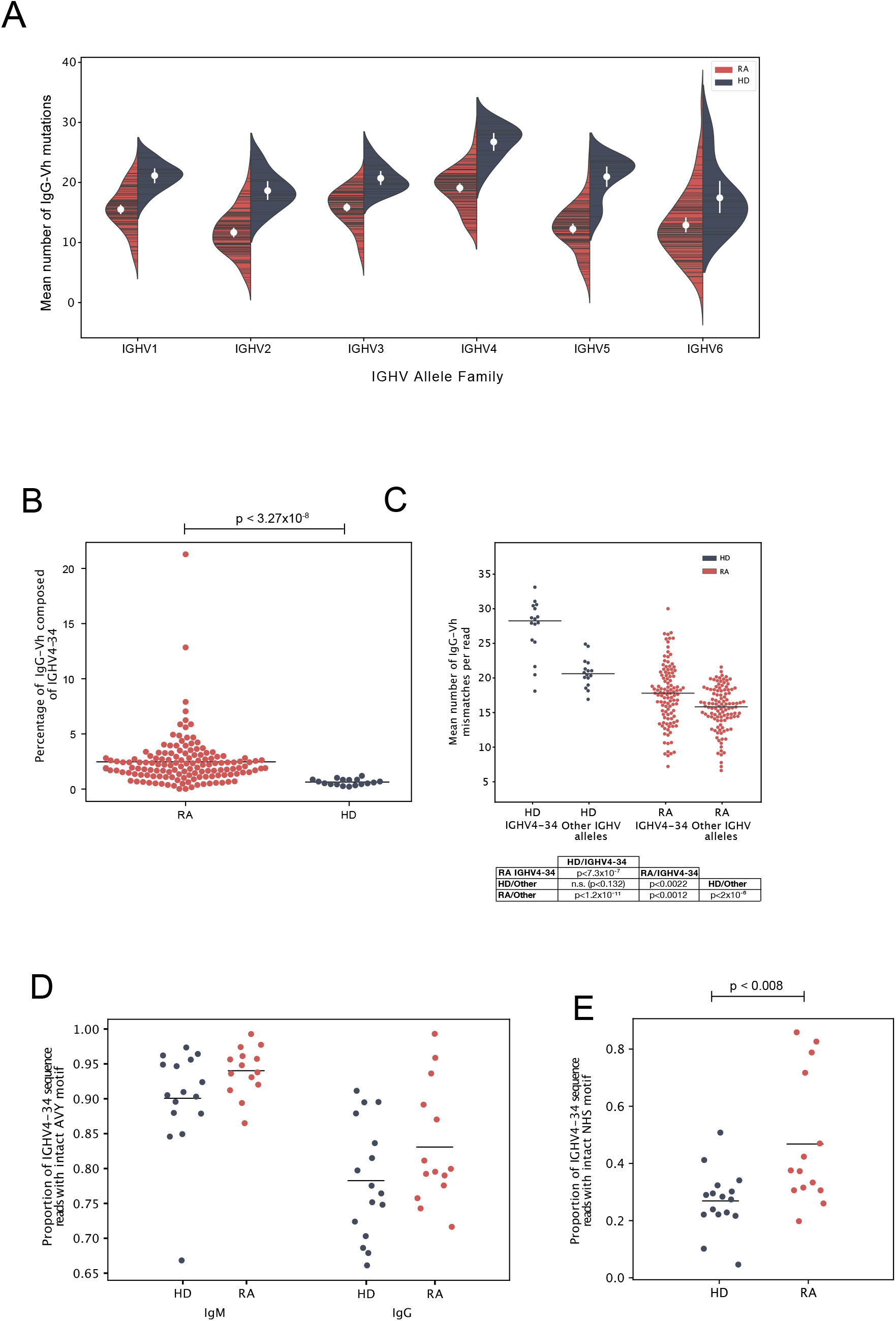
A. The mean number of IgG-Vh V segment mismatches per read for each individual in the ERA (n=113) and healthy control groups (n=16). Data are split by germline IGHV family group. White circles denote group means, vertical white lines show the 95% confidence interval for the mean. B. Percentage of IgG reads that use the IGHV4-34 allele in ERA patients (Cohort 2, n=113) and control donors (cohort 1, n=16). P value calculated using Kruskal Wallis with Dunn’s post-hoc pairwise test, and with Holm-Šídák correction for multiple comparisons of group means. C. Mean number of IgG-Vh mismatches per read for ERA donors (n=113, cohort 2) and heathy control donors (n=16). For each donor, the mean number of mutations for all reads mapping to IGHV4-34, or to other IGHV alleles, were calculated and plotted independently. D. Proportion of IGHV4-34 reads of IgG and IgM Isotypes where the carbohydrate binding AVY motif within framework region 1 (IMGT numbering 24-26) has an unchanged amino acid sequence. E. Proportion of IGHV4-34 IgG sequences where the NHS glycosylation motif within CDR2 (amino acid residues 57-59, IMGT numbering scheme) has an unchanged amino acid sequence. P value calculated using t-test (2 tailed) to compare the group means.

### IgG^hypoM^ sequences are not caused by oligoclonal expansions of poorly mutated IgG+ve BCRs

To confirm the polyclonal nature of the IgG^hypoM^ BCR sequences, an equality metric called the Gini coefficient was used to compare the degree of clonal expansion in the hypo- and hyper-mutated compartments of each patient’s repertoire. The Gini coefficients of the IgG^hypoM^ and hypermutated IgG repertoires of each patient from cohort 2 were similar, indicating that both the hypo- and hyper-mutated components of the repertoire have similar clonotypic frequency structures [Figure 3A]. We further investigated the degree of clonal dominance of IgG^hypoM^ compared to hypermutated sequences from RA donors at the time of diagnosis and healthy controls derived from cohort 1 where an average of 200,000 BCRs were sequenced per donor. Concurring with two recently published reports on NGS of RA BCRs, the majority of RA patients and healthy controls expressed some clonotype frequencies greater than 0.5% of the total sequence reads(26, 27) [Figure 3B]. Within these repertoires, both hypermutated and IgG^hypoM^ sequences exhibited a similar distribution of clonotype frequencies, including a very large number of clonotypes with single reads [Figure 3C]. One explanation for the higher frequency of IgG^hypoM^ in RA donors could be the failure of enzymes involved in somatic hypermutation (SHM), such as activation-induced cytidine deaminase (AID) or the subsequent mismatch repair enzymes. However mutations were preferentially targeted to the same regions of the IgG-Vh segment in RA and control donors, and there were no inter-group differences in the targeting of mutation hotspots [Supplementary Figures 4-5]. Taken together, these results indicate that the increased prevalence of IgG^hypoM^ in RA donors did not result from AID or mismatch repair enzyme impairment or to the mono- or oligo-clonal expansions of IgG+ve BCRs with few mutations.

**Figure 3.**
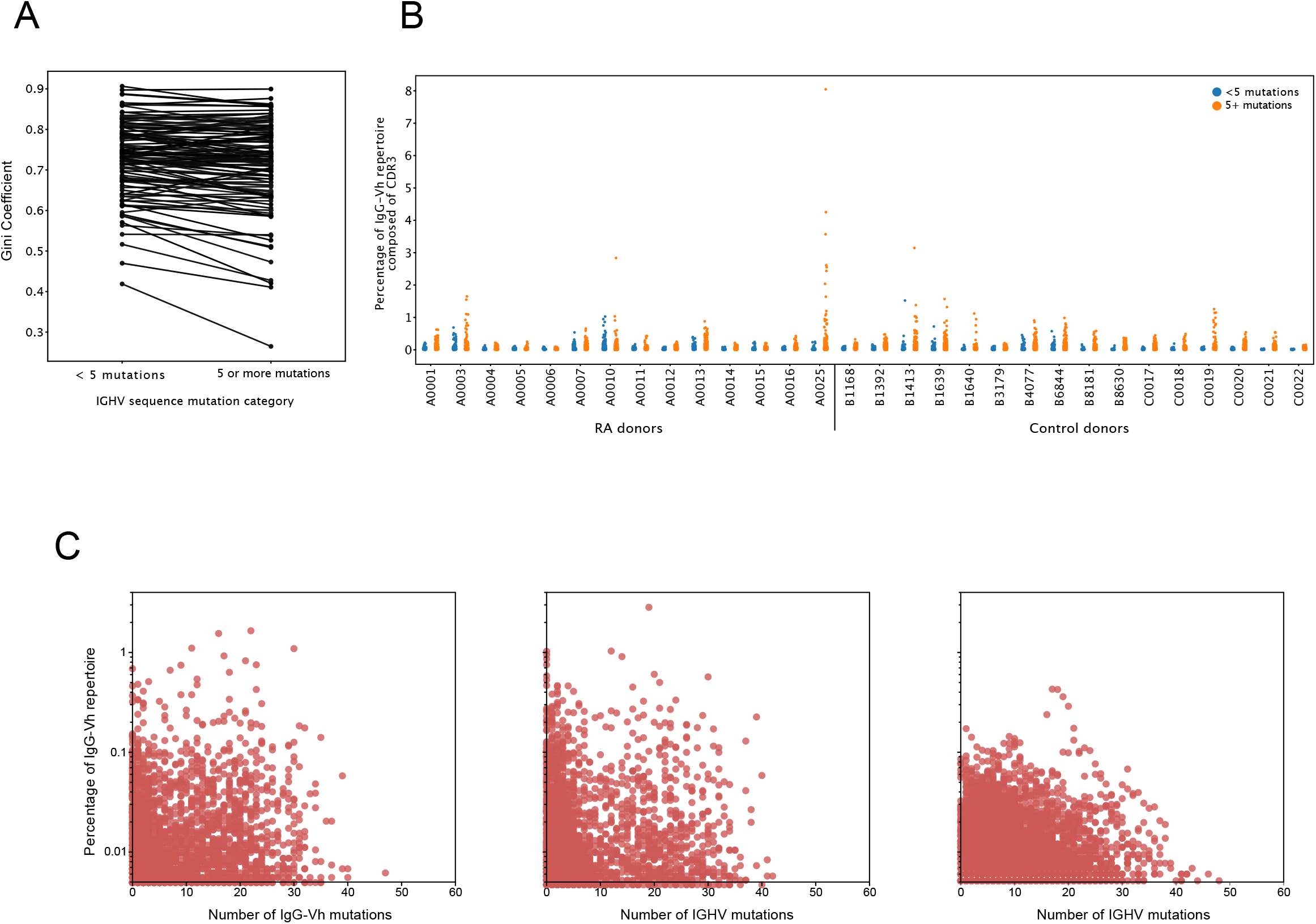
A. Gini coefficients of IgG sequences for each RA donor from cohort 2 (n=113). Gini coefficients are a measure of inequality of distribution, where a value of 0 indicates perfect equality (all IgG clonotypes of equal proportion). The Gini coefficient was calculated independently for hypomutated (Fewer than 5 mismatches) or hypermutated (5 or more mismatches) sequences to compare the degree of clonal expansion in each category. B. Percentage of the IgG-Vh repertoire composed of unique CDR3 clonotypes from ERA patients (cohort 1, n=14), with sequences split into hypermutated (5 mismatches or more) and hypomutated (fewer than 5 mismatches). C. As for 3B from the three individual RA donors with the greatest frequency of hypomutated B-cells (in cohort 1).

### The BCR^hypoM^ are expressed by IgG+CD27-ve B cells

The proportion of IgG^hypoM^ was significantly higher in the IgG^+ve^CD27^−ve^ B-cell population from RA patients than from either the IgG^+ve^CD27^+ve^ RA population or from the same population in the healthy controls [Figure 4A, Mann-Whitney U, p<0.015]. The absolute number of circulating peripheral blood double negative (IgM^−ve^IgD^−ve^CD27^−ve^) B cells in RA patients was also significantly increased at the time of diagnosis and did not change following six months therapy [Figure 4B]. Within individual RA patients, the number of double negative B cells at baseline and following six months of DMARD therapy was still clearly correlated, suggesting that they did not decrease significantly with synthetic DMARD treatment [Figure 4C]. Importantly the increase in double negative B cells was also reflected in a significant increase in the frequency of IgG^+ve^CD27^−ve^ B cells [Figure 4D]. IgG^+ve^CD27^−ve^ B-cells from RA patients expressed significantly less CD24, CD21 and CD138 but similar levels of CD38, CD73 and CD1c when compared to the same subset in healthy controls [Figure 4E].

**Figure 4.**
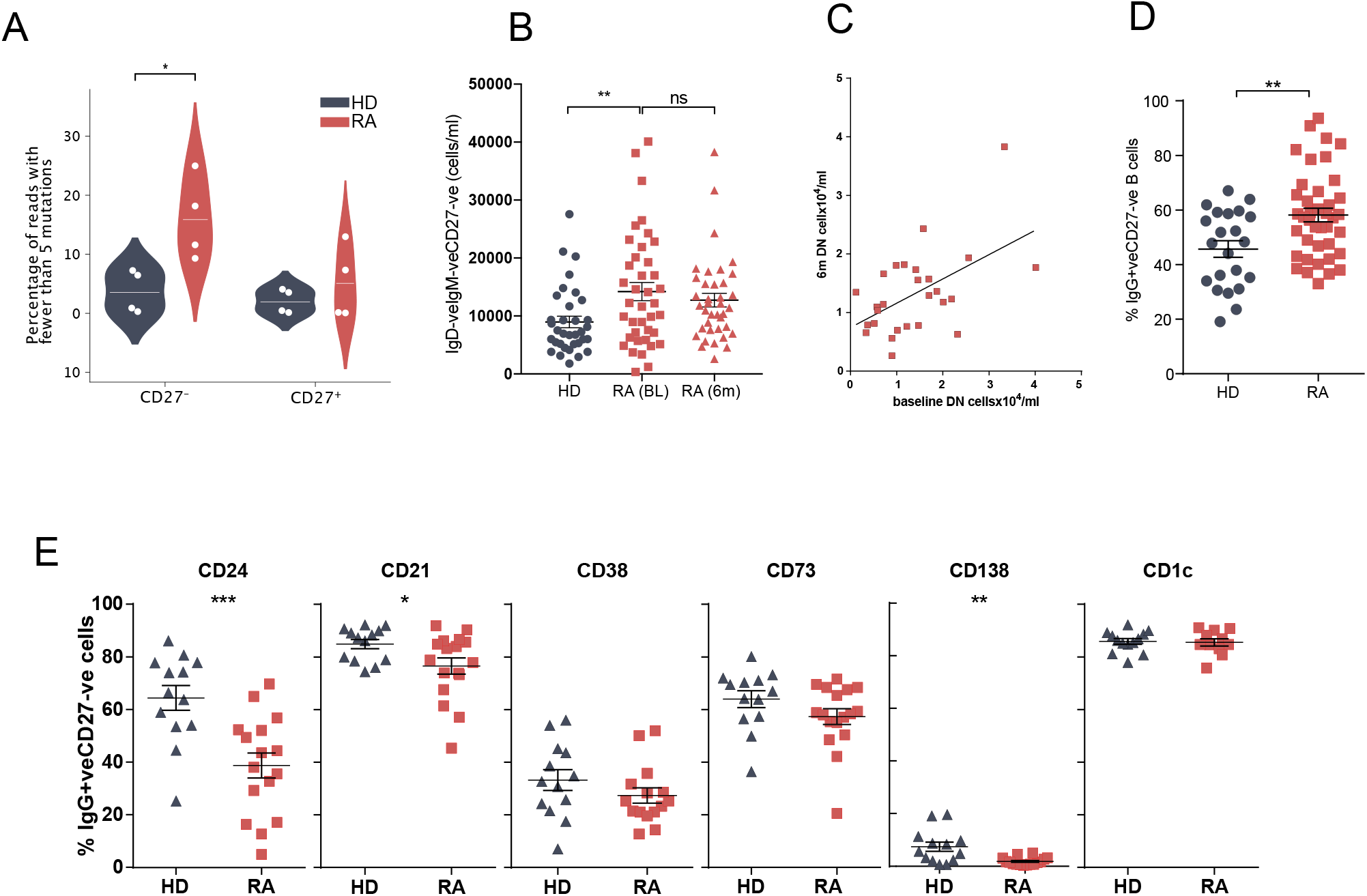
A. Prevalence of hypomutated V segment sequences in CD27^+ve^ and CD27^−ve^ IgG^+ve^ B-cells from RA and control donors. White spots indicate individual data points for each donor (n=4 per group) B. Whole blood was stained for flow cytometry to determine cell number. IgD^−ve^IgM^−ve^CD27^−ve^ B cells were gated and cells/ml calculated. N=35 HD, 40 RA(0m), 39 RA(6m). C. Paired baseline and 6 month data from Fig 4B was used to determine if the frequency of IgD^−ve^IgM^−ve^CD27^−ve^ B cells was correlated in the same patient 6 months following DMARD therapy. D. PBMCs from control donors and RA patients were stained for flow cytometry. Cells were gated for CD20^+ve^CD19^+ve^IgG^+ve^ B cells then the proportion of CD27^−ve^ cells determined. N=22 HD, 40 RA. E. Cell surface markers of IgG^+ve^CD27^−ve^ B cell population were analysed by flow cytometry. Cells were gated as Fig 4D. N=13 HD & 16 RA

### IgG^+ve^CD27^−ve^ B cells are enriched in the synovium and secrete pro-inflammatory cytokines

Utilising paired blood and synovial tissue from ESRA patients undergoing joint arthroplasty (cohort 4/ Supplementary table 4), revealed that the synovium was enriched for both IgG^+ve^ B cells [Figure 5A] and IgG^+ve^CD27^−ve^ B cells [Figure 5B]. Compared to circulating IgG^+ve^CD27^−ve^ B cells, the expression of FcRL4, RANKL, CD73 and CD138 was upregulated, whilst HLADR, CD21, CD40 and CD38 were expressed on fewer synovial IgG^+ve^CD27^−ve^ B cells [Figure 5C]. Synovial IgG^+ve^CD27^−ve^ B-cells expressed significantly more TNF-alpha but similar amounts of GM-CSF than either IgG^−ve^CD27^−ve^ naïve or IgG^+ve^CD27^+ve^ memory B cells of patients with established RA [Figure 5D].

**Figure 5.**
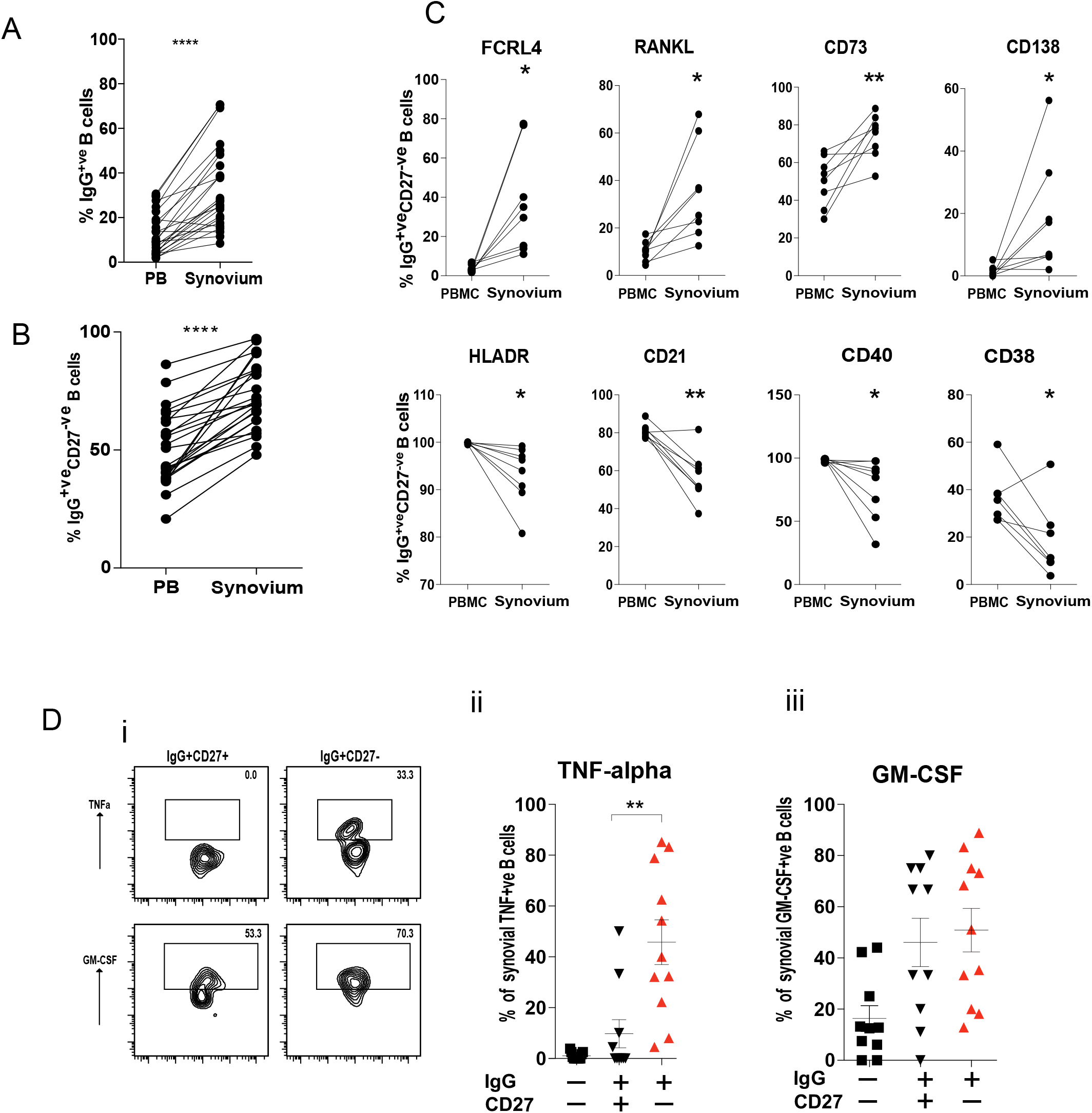
A. Flow cytometry of paired peripheral blood (PB) and synovial tissue B cells from RA patients taken at the time of undergoing arthroplasty. The graph shows the percentage of IgG^+ve^ B cells within the CD20^+ve^CD19^+ve^ B cell population. N=24 B. CD20^+ve^ CD19^+ve^ population as for (A). Graph shows the percentage of IgG^+ve^ CD27^−ve^ B cells within this B cell population. N=24 C. Plots show the percentage of cells positive for the surface markers of paired PB and synovial samples within the IgG^+ve^ CD27^−ve^ B cell subset. N=8 D. Representative flow cytometry plots showing the intracellular cytokine staining of stimulated IgG^+ve^ CD27^−ve^ and IgG^+ve^ CD27^+ve^ B cell subsets for TNF-alpha and GM-CSF (i). Pooled data for (ii) TNF-alpha and (iii) GM-CSF. Each point represents an individual patient sample. N=11

### Peripheral blood and synovial B cell repertoires are distinct in RA

As well as secreting cytokines, synovial B cells are reported to secrete autoantibodies that have undergone SHM within GC like structures(26, 28–30). Paired peripheral and synovial IgG-Vh sequences from RA patients undergoing arthroplasty were sequenced [Figure 6A]. To avoid any potential confounding that may occur with increased levels of mRNA in plasma cells, purified CD20^+ve^ B cells were analysed. In six out of eight patients the lower quartile of the distribution of somatic mutations was higher in the periphery than in the synovium. The mean number of mutations displayed by the peripheral B cell repertoire of patients (mean 15.27 ± 5.09) was lower than the mean number of mutations displayed by the paired synovial repertoires (mean 19.65 ± 2.96, paired t-test, p=0.02). The RA synovium has been reported to harbour clonal expansions of B cells(26, 30). The percentage of the repertoire composed of each unique CDR3 clonotype, was plotted, dividing the BCR sequences into those with greater or fewer than five mutations relative to the predicted germ line sequence [Figure 6B]. No clear preference was noted within either the hypermutated or IgG^hypoM^ sequences in terms of CDR3 clonal frequencies with all four populations expressing CDR3 clonal frequencies greater than 0.5% of the IgG^+ve^ B cell repertoire.

**Figure 6.**
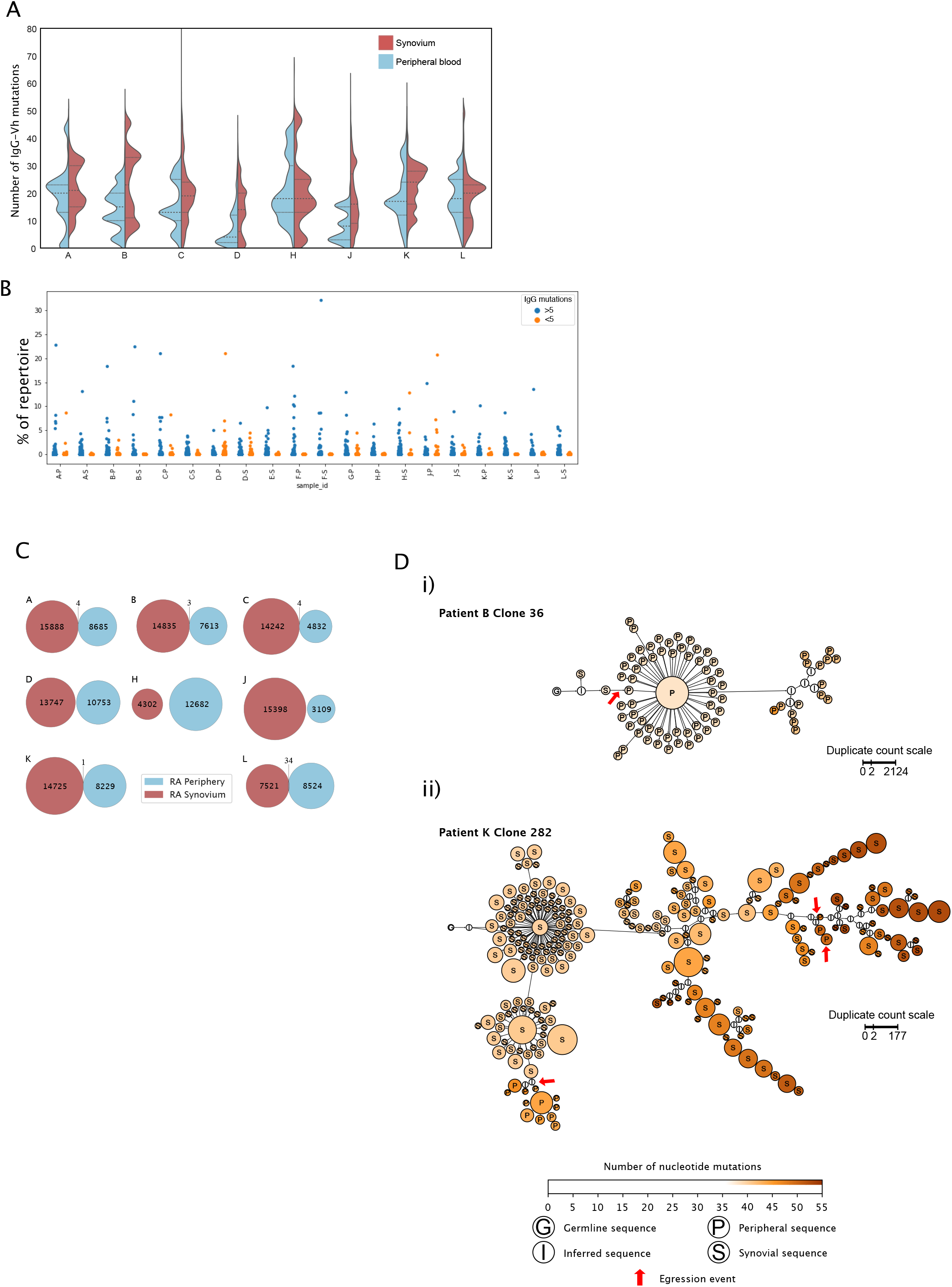
A. Distributions of mutation counts from paired peripheral blood (blue) and synovial (red) IgG sequence repertoires from each of 8 RA patients with ESRA undergoing arthroplasty. The central horizontal line indicates the median of each distribution, with the upper and lower dashed lines representing the upper and lower quartiles respectively. B. Percentage of the repertoire composed of each CDR3 clonotype for paired synovial (S) and peripheral blood (P) samples from each of the 8 RA patients shown in 6A (cohort 4). C. Repertoire overlap of synovial (red) and peripheral blood (blue) IgG repertoires of the RA patients. Each Venn diagram represents a single patient. The number of unique, non-singeton IgG sequences in the repertoire from each compartment is depicted at the centre of each circle, and shared sequences are enumerated at the intersection between the two circles. Two shared sequences are considered identical if they possessed the same CDR3 nucleotide sequence and used the same V and J gene segments. D. Lineage trees of B cell clones that show evidence of egression from the synovium into the periphery of RA patients, inferred for i) clone 36 from patient B and ii) clone 282 from patient K. Each node represents a unique non-singleton IgG sequence with the size of the node scales non-linearly in proportion to the number of sequence duplicated observed. The label at the centre of each node represents the tissue origin of the sequence, and node colour indicates the number of somatic mutations present in the clone sequence. Red arrows mark egression events from the synovium to the periphery. Lineage trees were inferred using PHYLIP v3.6 and plotted in Gephi v0.92.

Finally it was important to determine if RA patients shared BCR sequences that may point to particular pathogenic clonal expansions. The term ‘public sequence’ is used to describe similar or identical T cell receptor (TCR) or BCR sequences which may arise in different individuals, indicative of a convergent immune response in different individuals to a common antigenic stimulus(31). This may have particular relevance for autoimmune mediated tissue damage including RA. We hypothesised that identical CDR3 sequences would be found in more than one patient either in newly diagnosed DMARD naïve RA patients (cohort 1) or within the paired blood and synovial samples of the ESRA patients (cohort 4). We also compared these CDR3 sequences to healthy subjects from the cohort 1 study. Notably, no common CDR3 sequences were found to be shared by the peripheral IgG sequences of the healthy subjects and only 21 CDR3 sequences were shared (ie present in at least 2 individuals) by the peripheral B cell repertoires of RA patients, comprising a tiny fraction of the total repertoire. In addition very few CDR3 sequences were present in both the blood and synovium of the same patient [Figure 6C]. Where overlap was seen (in the repertoires of 5 RA patients), it made up less than 0.1% of the synovial repertoire. To confirm this clonal lineage analysis of all shared sequences was employed to detect any evidence of egression events from the synovial compartment. Only two lineage trees, from patient B and patient K, showed evidence of egression events from the synovium to the periphery [Figure 6D]. This shows that shared CDR3 sequences were not abundant in the peripheral B cell repertoires of RA patients and that synovial and peripheral blood repertoires are largely independent. In addition, given the extremely low support for egression events from the synovium, there is no evidence to support the hypothesis that the IgG^hypoM^ sequences in the peripheral blood B cell repertoires arise in the inflamed synovial joint.

## Discussion

This is the largest study to date, utilising NGS, to examine the IgG-Vh repertoires of over 150 RA patients. RA patients express significantly more hypomutated B cell receptors within IgG^+ve^CD27^−ve^CD24^lo^CD21^lo^. In patients with established RA, IgG^+ve^CD27^−ve^CD24^lo^CD21^lo^ B cells are enriched in the synovium, where they upregulate the expression of FcRL4, RANKL, CD73, CD138 and secrete TNF. Significantly more IgG-Vh utilise the auto reactive allele IGHV4-34, which are again distinctively less mutated than the same allele expressed by healthy controls. Surprisingly there is virtually no sharing of the repertoire between the blood and synovium or between RA patients. The accumulation of IgG^hypoM^ expressing CD27^−ve^CD24^lo^CD21^lo^ B cells may both augment the formation of self-reactive immune complexes and pro-inflammatory cytokines. We hypothesise that following B cell depletion therapy, the frequency of IgG^hypoM^ would gradually increase to a critical threshold prior to a flare of synovitis.

A previous report alludes to the loss of clonally expanded populations of B cells from the peripheral blood into the synovium at the time of RA onset (27). Assessing a greater number of IgG sequences we observed expanded populations of B cells in both RA and healthy controls, making it difficult to see how dominant BCR clones could predict the onset of RA in at-risk individuals. Despite RA being a heterogenous disease, we expected to find particular IgGVh-CDR3 clones that were shared in the blood or synovium of RA patients. Instead the repertoire overlap analysis demonstrated a very low number of multi-compartmental IgG sequences when compared to the overall size of either repertoire, suggesting that the peripheral and synovial B cell repertoires are quite distinct within and between patients. The cause for this may, as previously reported, arise from the migration of peripheral blood B cells into the synovium(27). As synovial B cells express distinct chemokine receptors from peripheral B cells, they may be sequestered within the synovium and rarely be observed in the peripheral blood again(32). In line with this, we observed only two B cell clones that showed evidence of a migration event from the synovium into the peripheral blood. We also noted a scarcity of public CDR3 sequences in the peripheral B cell repertoire between RA patients, which was surprising if RA is an antigen-driven disease. Given that different immunoglobulins with similar (but not identical) physiochemical properties can recognise the same antigen epitope(33), it is still possible that IgG sequences that react to the same epitope are not identical.

The stimuli driving the emergence of such a high frequency of class switched IgG^hypoM^, that approached 40% of the IgG-Vh repertoire in some RA patients, is currently unknown. Central and peripheral tolerance checkpoints are known to be defective in RA patients, leading to the accumulation of naïve autoreactive B cells in the periphery(34). Activation of naïve B cells, facilitated by T cell help and/or TLR ligands, induces class switching outwith the germinal centre(35). The high frequency of IgG^hypoM^ in RA patients stongly suggests that they have arisen from an ongoing extrafollicular response. Given the lack of NHS N-glycosylation motif mutations within the self-reactive IGHV4-34 IgG sequences, we hypothesise that these IgG^hypoM^ B cells arise from an expanded pool of self-reactive innate like B cells(25). CD27^−ve^IgD^−ve^ B cells are increased in the blood of RA patients both at diagnosis and following synthetic DMARD therapy(36–38). CD27 is widely used as a marker of human memory B cells, but CD27^−ve^IgG^+ve^ B cells also contain a pool of short-lived memory cells(8, 39, 40). They are substantially increased in the blood of patients with SLE, correlate with a higher disease activity index(41) and are significantly diminished in SLE patients following rituximab therapy(42). Over time an increasing number of RA patients are failing to respond to multiple synthetic and biologic DMARD therapies. Future work will explore if these refractory RA patients express more IgG^hypoM^ CD27^−ve^IgG^+ve^ B cells and if their emergence following BCDT foretells clinical disease relapse.

## Abbreviations

BCR: B cell receptor
DMARD: Disease modifying anti-rheumatic drug
V_H_: variable portion of the heavy chain
BCDT: B cell depletion therapy
ACPA: anti citrullinated protein antibodies RF-rheumatoid factor
NGS: next generation sequencing
PBMC: peripheral blood mononuclear cells
ERA: early rheumatoid arthritis
ESRA: established RA
CDR: complementarity determining region
SHM: somatic hypermutation
AID: activation-induced cytidine deaminase
GC: germinal centre
FR: framework region
CS: class-switching

## Author contributions

GC, KM and CG carried out experiments. GC, LC, SG undertook the bioinformatics analysis. HJ and SB contributed to clinical data and/or sample collection. MG, GC and DG designed the experiments, analyzed the data, and wrote the manuscript. IMcI and HJ reviewed the manuscript and IMcI managed the SERA inception cohort.

## Acknowledgements

This work has made use of the resources provided by the Edinburgh Compute and Data Facility (ECDF) (http://www.ecdf.ed.ac.uk/). The authors thank the staff from the QMRI Flow Cytometry Facility for help with cell sorting.

