## Supplementary Figures for "Rheumatoid arthritis patients express a skewed repertoire of polyclonal, hypomutated B-cell receptors"

Supp Figure 1

**A**

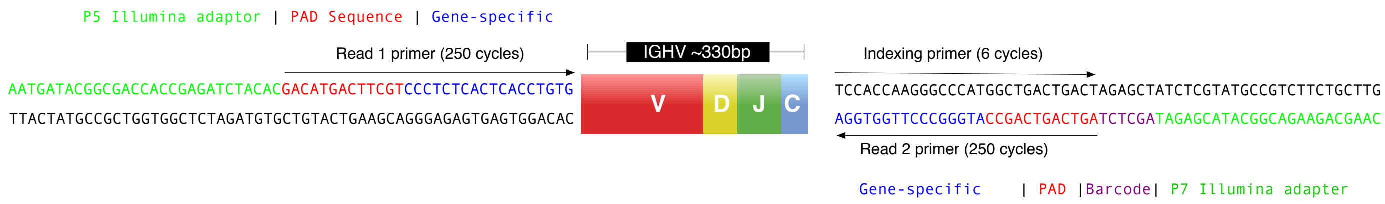


**B**

| | |
|---|---|
| **IGHV FR1 forward amplification primers** | |
| hIGHV2.0 | AATGATACGGCGACCACCGAGATCTACACGACATGACTTCGTCCCTCTCACTCACCTGTG |
| hIGHV2.1 | AATGATACGGCGACCACCGAGATCTACACGACATGACTTCGTCCCTKAGACTCTCCTGTG |
| hIGHV2.2 | AATGATACGGCGACCACCGAGATCTACACGACATGACTTCGTCCCTGARACTCTCCTGTG |
| hIGHV3.0 | AATGATACGGCGACCACCGAGATCTACACGACACTACAGCGTAGACCCTCACRCTGACCT |
| hIGHV0.0 | AATGATACGGCGACCACCGAGATCTACACGACATAGACCGGTTCAGTGAAGGTYTCCTGC |
| hIGHV1.0 | AATGATACGGCGACCACCGAGATCTACACCGCCGGAAGCAGTGCTGAGGTGAAGAAGCCT |
| hIGHV1.1 | AATGATACGGCGACCACCGAGATCTACACCGCCGGAAGCAGTYCAGGACTGGTGAAGCCT |
| hIGHV1.2 | AATGATACGGCGACCACCGAGATCTACACCGCCGGAAGCAGTSCAGGACTGTTGAAGCCT |
| hIGHV4.0 | AATGATACGGCGACCACCGAGATCTACACCATGTGCCTGTGTGTCCCTGAGACTCTCCTG |
| hIGHV4.1 | AATGATACGGCGACCACCGAGATCTACACCATGTGCCTGTGTGTCTCTGARGATCTCCTG |
| hIGHV5.0 | AATGATACGGCGACCACCGAGATCTACACCTTAGAGTCACGTCTCCTGCAAGGYTTCTGG |
| **IgGHC reverse primer** | |
| hIGGHCrev(INDEX1) | CAAGCAGAAGACGGCATACGAGATAGCTCTAGTCAGTCAGCCATGGGCCCTTGGTGG*A |

Supplementary Figure 1

A: PCR amplicon sequence and sequencing strategy. Non-template encoded sequences are incorporated into amplicons during PCR as shown. Read 1 primer, Indexing primer and Read 2 primer locations are indicated by the black arrows.

B: Primer sequences for amplification of IgG and IgM repertoires. Primers contain a hexameric indexing sequence, for illustration only a single index is shown for each reverse primer.

Supplementary Figure 2

A

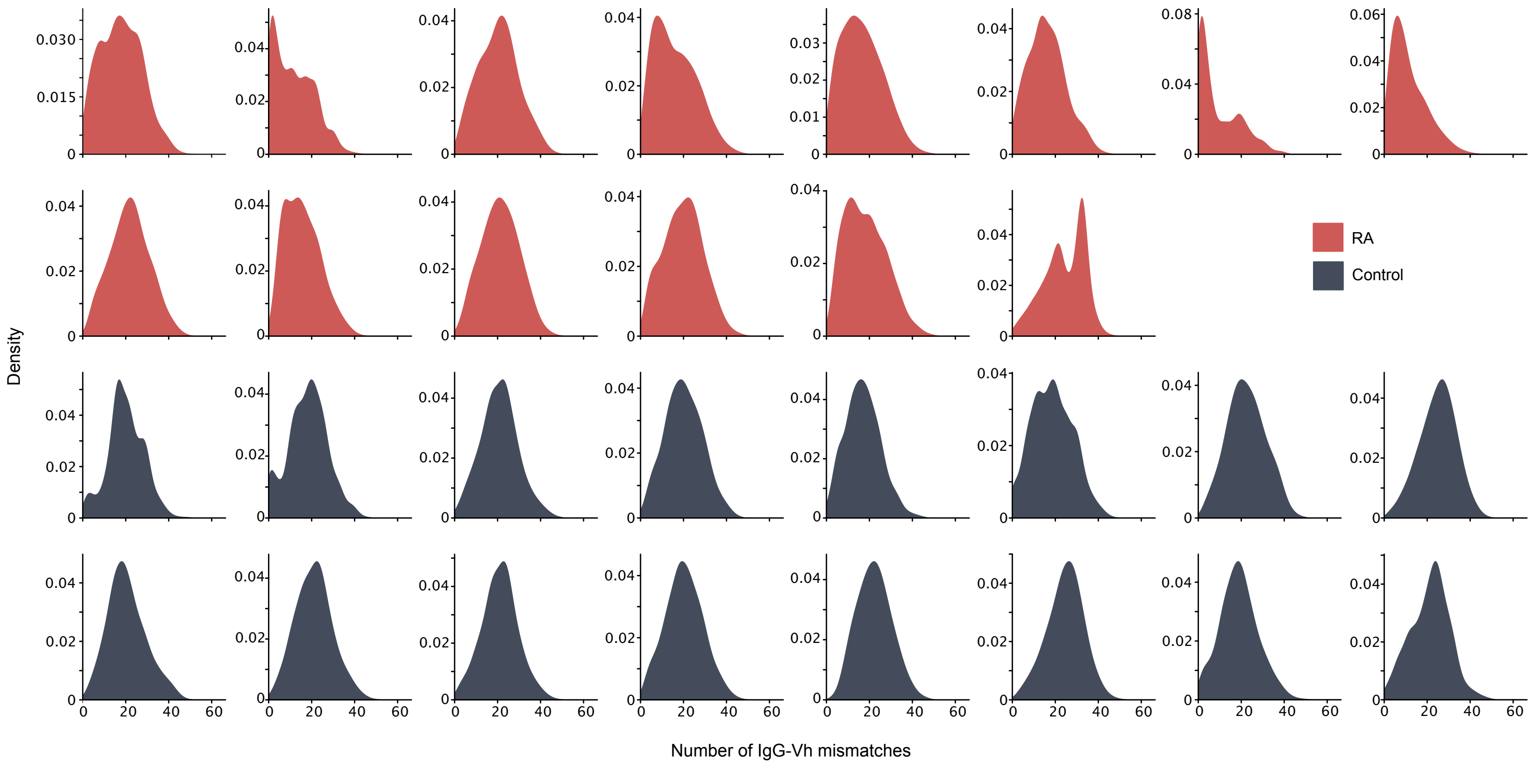


B

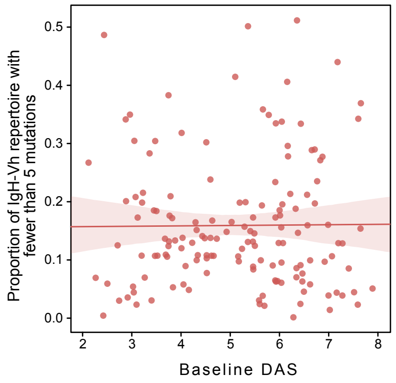


<u>Supplementary Figure 2</u>

A: Distribution of the number of IgG-Vh mismatches for each sequencing read of ERA (n=14) and control donors (n=16) from cohort 1.

B: Linear regression of the proportion of the IgG-Vh reads from each repertoire against the disease activity score (DAS) at the time of diagnosis in cohort 2. Spearman correlation coefficient = -0.024832. The regression line is show as a solid red line, with the 95% confidence interval in indicated by the surrounding shaded area.

Supp Figure 3

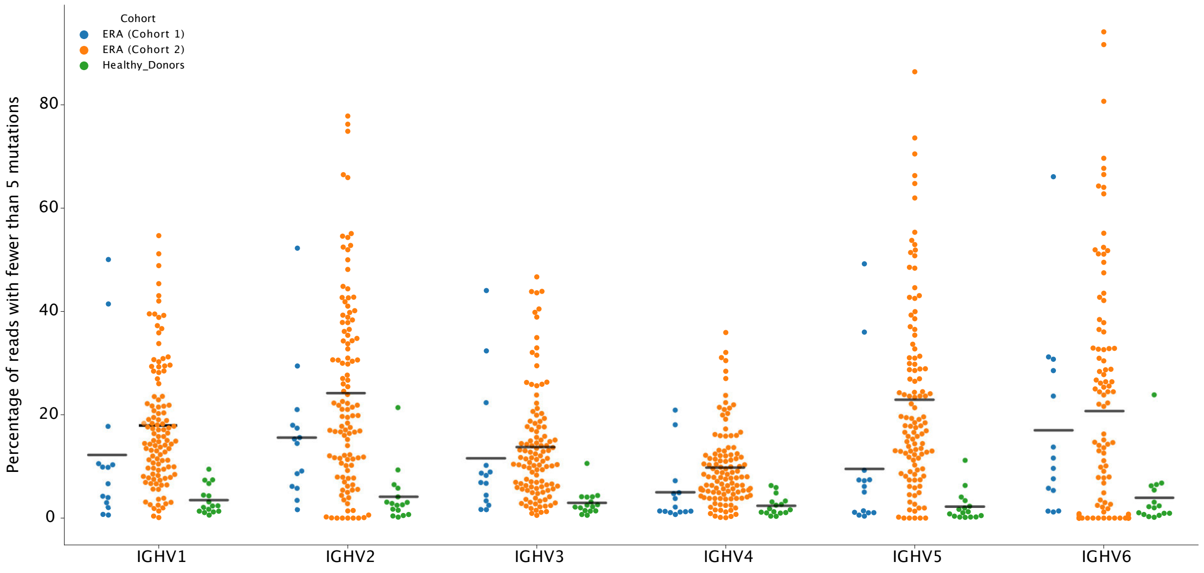


Supplementary Figure 3

Percentage of IgG reads from ERA and healthy donors that have fewer than 5 mutations, split by germline IGHV allele family. Only reads that map to the six most frequent IGHV families (IGHV1-6) are shown to ensure sufficient reads are available in each donor to be representative. Hortizontal bars denote the group mean.

Supp Fig 4

i) Early RA donors

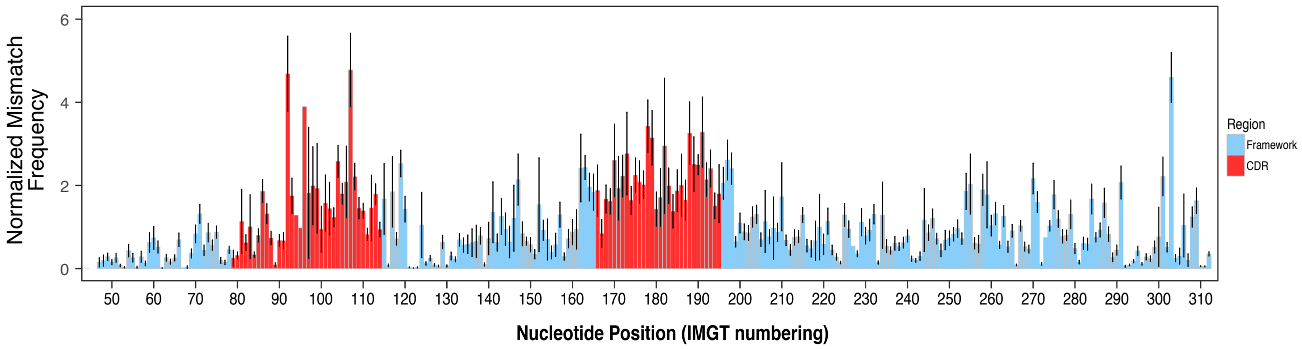


ii) Healthy control  donors

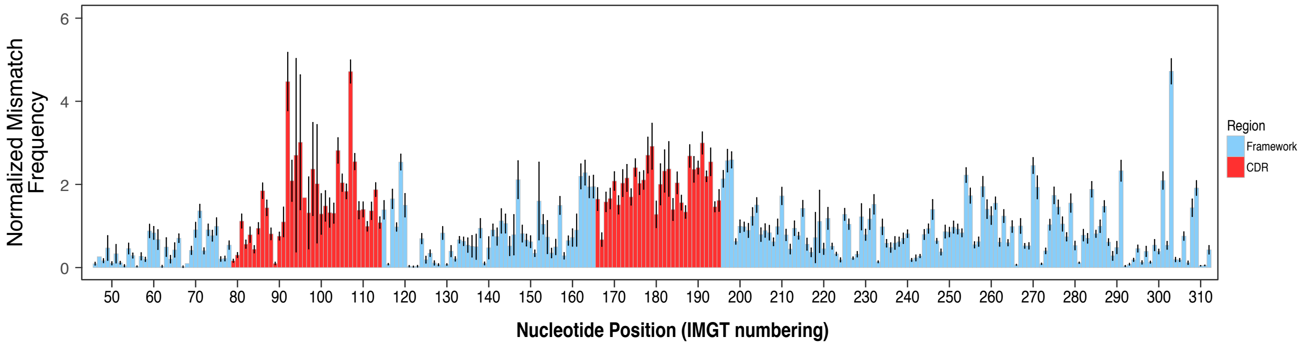


<u>Supplementary Figure 4</u>

Average frequency of mismatches along the V segment for IgG sequences i) RA and ii) healthy control donors form cohort 1 (n=14 & n=16 respectively). The normalised mismatch frequency is calculated as the number of times a mismatch appears at the IMGT position divided by the number of times the IMGT position appears within the aligned sequences of each patient. The group mean is calculated from individual patient frequencies at each IMGT position; error bars show 95% CI of the mean. Mean and 95% CI are normalized by the group mean for per-base mismatch frequency. Red bars show the CDR and blue bars show the FR regions.

### Supplementary Figure 5

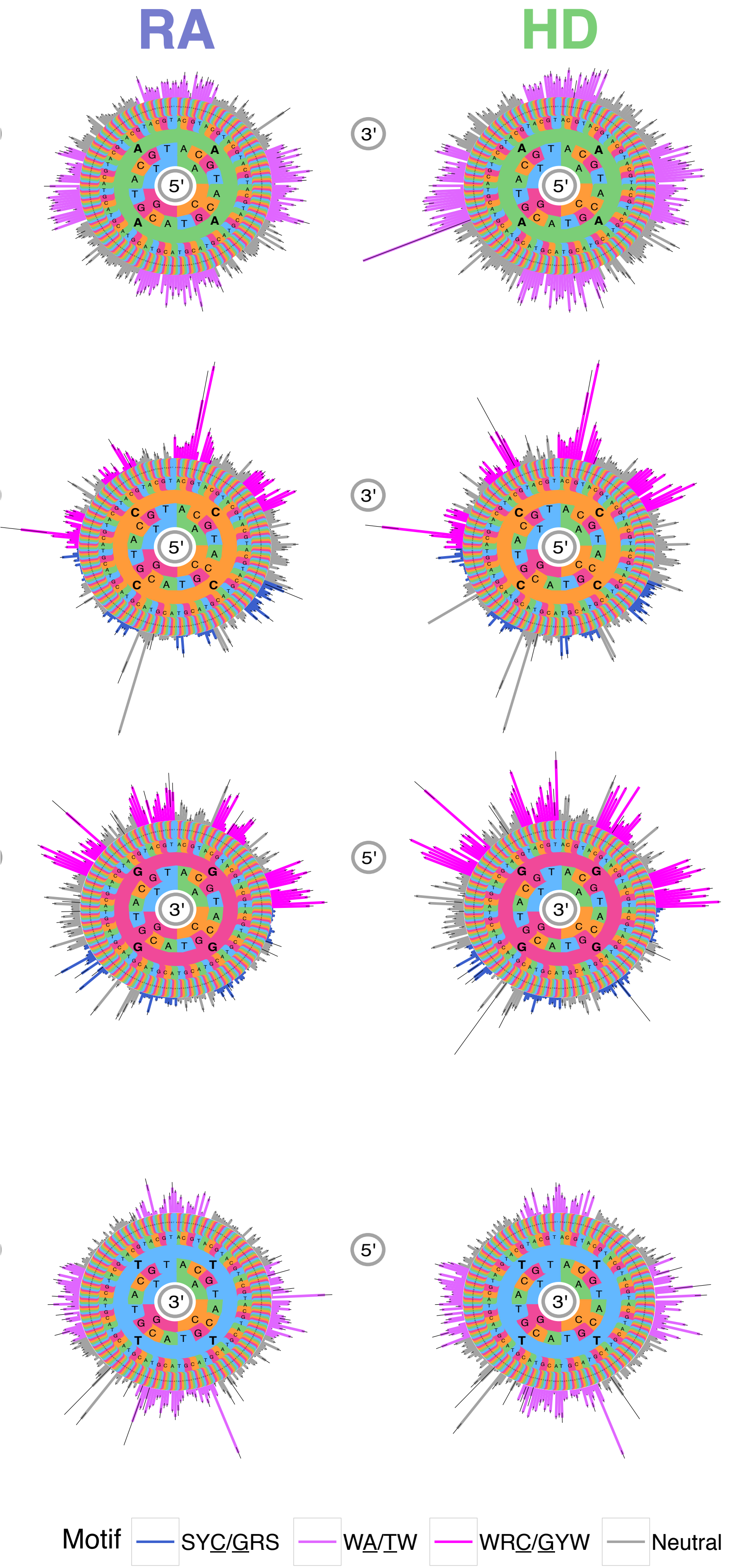


Motif Mutabilities of All Mutations in IgG repertoires of individuals from cohort 1 (n=16 for HD and n=14 for RA). Analysis of motif mutability in all mutations to assess targeting by SHM and selection. Data are shown as hedgehog plots: Circular bar graphs show the group mutation frequency of the central nucleotide in a 5-nucleotide kmer. The central set of concentric circles shows the initial sequence of the 5-mer and the colour of the bars shows the motif identity (hotspots, coldspots, and neutral). Mutation frequency is calculated as the number of times the central nucleotide is mutated, divided by the number of times the kmer appears in a patient. Group frequencies are calculated as the mean of each patient kmer mutation frequency and normalized by the per-base mismatch frequency for that group. Error bars show the 95% CI of the mean. R = (A or G), S = (C or G), Y = (C or T), and W = (A or T)

Supplementary Table 1

Cohort 1: Newly diagnosed RA and healthy controls.

| | Cohort 1 ERA (n=14) | Cohort 1 HD (n=16) |
|---|---|---|
| Age (year, IQR) | 52.6 (36;66.5) | 53.1 (43.5;60.3) |
| Female (number) | 11 (78.6%) | 10 (62.5%) |
| CCP (IU/ml, IQR) | 290.8 (94;503.3) | N/A |
| RF (IU/ml, IQR) *1 | 86.4 (15.9;67.9) | N/A |
| Baseline DAS28 (IQR) *2 | 5.4 (4.8;6.4) | N/A |
| Follow up DAS28 at 6 months (IQR) *3 | 4.1 (2.8;5.6) | N/A |

*1n=9, *2n=10, *3n=9

Supplementary Table 2

Cohort 2:Scottish Early Arthritis (SERA) study

(i) Baseline data

| | SERA study (n=113) |
|---|---|
| Age (year, IQR) | 61.9 (53.6;71.1) |
| Female (number) | 65 (57.5%) |
| CCP (IU/ml, IQR) *1 | 210.2 (5.7;340) |
| RF (IU/ml, IQR) *2 | 175.2 (15;348.5) |
| Baseline DAS28 (IQR) | 5.0 (3.8;6.1) |
| Follow up DAS28 at 6 months (IQR) | 3.8 (2.3;5.3) |

*1n=97, *2n=64

(ii) SERA: 6 months post DMARD treatment

| | SERA study (n=12) |
|---|---|
| Age (year, IQR) | 64.6 (54.9;79.7) |
| Female (number) | 8 (66.7%) |
| CCP (IU/ml, IQR) | 186.9 (38;340) |
| RF (IU/ml, IQR)*1 | 231 (20;601) |
| Baseline DAS28 (IQR) | 5.1 (3.6;6.4) |
| 6 month DAS28 (IQR) | 4.9 (3.3;6.7) |
| DMARD treatment | |
| − Methotrexate | 10 |
| − Hydroxychloroquine | 4 |
| − Sulfasalazine | 5 |
| − Leflunomide | 1 |
| Number of concurrent DMARDS | |
| − One | 7 |
| − Two | 2 |
| − Three | 3 |

*1n=3

Supplementary Table 3

Cohort 3:Patients with established RA

that had failed synthetic DMARD therapy

| | Cohort 3 N=16 |
|---|---|
| Age (years, IQR) | 52.6 (37;67.8) |
| Gender (female) | 12 (75%) |
| CCP (IU/ml, IQR)*1 | 232.1 (61.8;405) |
| RF (IU/ml, IQR) *2 | 69.3 (10.6;131) |
| DMARD (current/previously failed) | |
| − Methotrexate | 16 |
| − Hydroxychloroquine | 12 |
| − Sulfasalazine | 15 |
| − Leflunomide | 10 |
| − Gold | 2 |
| − Cyclosporin | 1 |
| Number of previous Biologic DMARDs | |
| − None | 1 |
| − One | 6 |
| − Two | 6 |
| − Three | 2 |
| − Four | 1 |
| Biologic agents (current/previously failed) | |
| − Certolizumab | 14 |
| − Etancercept | 6 |
| − Adalimumab | 1 |
| − Rituximab | 4 |
| − Tocilizumab | 2 |
| − Abatacept | 1 |

*1N=14, *2N=11

Supplementary Table 4

**Supplementary table 4: Methods summary for cell preparation and B cell repertoire amplification**

| Cohort | Description | Cell site and RNA preparation | Amplification method | Isotype(s) amplified* |
|---|---|---|---|---|
| 1 | Early RA (ERA) and control donors, n=14 and n=16 | PBMC purification, CD19+MACS | PCR amplification using IGHV FR1 primers, IGHC primers | IgM & IgG |
| 2 | Early RA (ERA), n=113 | Paxgene tubes - 3mL whole blood. | PCR amplification using IGHV FR1 primers, IGHC primers | IgG |
| 3 | Established RA (ESRA), n=16 | PBMC purification, CD19+MACS | PCR amplification using IGHV FR1 primers, IGHC primers | IgG |
| 4 | CD27 sorted B cells | CD20+, IgG+, CD27+or CD27- | SMARTseq2 whole-transcriptome amplification followed by PCR amplification using IGHV FR1 primers, IGHC primers | IgG |

*for primer information, see supplementary figure 1

SupplementaryTable 5

| Supplementary Table 5: Antibody reagents for flow cytometry | |
|---|---|
| Antibody Reagent | Clone Reference |
| CD3-BUV395 | UCHT1 |
| CD19-PE/Cy5 | HIB19 |
| CD20-AF700 | 2H7 |
| CD21-BV421 | B-ly4 |
| CD27-APC | M-T271 |
| CD38-BV510 | HIT2 |
| CD73-BV605 | AD2 |
| CXCR3-PE | IC6 |
| GMCSF-PE/CF594 | BVD2-21C11 |
| CD5-PE/Cy7 | L17F12 |
| CD11C-BV711 | 3-9 |
| CD21-PE/Cy7 | Bu32 |
| CD21-PerCP/Cy5.5 | Bu32 |
| CD24-BV711 | ML5 |
| CD24-FITC | ML5 |
| CD38-BV605 | HIT2 |
| CD40-APC/Cy7 | 5C3 |
| CD86-BV711 | IT2.2 |
| CD95(FAS)-BV510 | DX2 |
| CD138-PE/Cy7 | MI15 |
| CD138-PerCP/Cy5.5 | DL-101 |
| CD254(RANKL)-PE | MIH24 |
| HLA-DR-APC/Cy7 | L243 |
| IgD-APC/Cy7 | IA6-2 |
| T-Bet-PE/Cy7 | 4B10 |
| TNFα-BV510 | Mab11 |
| CD1c-PerCP/eFluor710 | L161 |
| CD20-eFluor450 | 2H7 |
| CD307d(FcRL4)-PerCP/eFluor710 | 413D12 |
| IgG-FITC | IS11-3B2.2.3 |
| IgG-PE | IS11-3B2.2.3 |
