## Supplementary Methods for "Rheumatoid arthritis patients express a skewed repertoire of polyclonal, hypomutated B-cell receptors"

### Online Supplementary Methods

#### B cell repertoire sequencing

First strand cDNA synthesis was performed using Superscript III first-strand synthesis supermix (Invitrogen) or total cDNA synthesis kit (Applied Biosystems). For samples where low cell counts were included (CD27 sorted B-cells and synovial cells, as indicated in Supplementary table 6), whole transcriptome amplification was performed using Smartscribe Reverse Transcriptase and Advantage2 PCR kits (Takara BioEurope) according to the Smart-Seq2 protocol(43). For all samples, V-region amplicons were generated by PCR using Phusion Flash polymerase (ThermoFisher Scientific) with individual pools of forward primers within framework region 1 (FR1) designed to amplify all known V-region alleles, and a reverse primer within the IgG or IgM constant regions (Supplementary Figure 1). 250bp paired-end sequencing was performed on an Illumina MiSeq sequencer using a pool of read 1 sequencing primers, an indexing primer, and read 2 constant region amplification primers. Read 1 and read 2 sequencing primer sequences were identical to the pool of amplification primers detailed in Supplementary Figure 1 but omitting the Illumina adaptor sequence.

#### Immune repertoire analysis

Sequence read-pairs were combined using the Flash utility (44). Sequence data were processed using the VDJfasta utility(45) or using the pRESTO, Change-O, Alakazam, and SHazaM packages of the Immcantation adaptive immune repertoire analysis framework (46, 47). Frequency distributions for each donor were derived using the ggplot2(48) and plyr(49) packages of the R statistical package(50). The Gini Index was calculated for each sample based upon the read counts for each unique complementarity determining region 3 (CDR3) amino acid sequence in the repertoire, according to the formula:

$$Gini\ index = \frac{\{2\sum_i^n i \times y_i\}}{\{n\sum_i^n y_i\}} - \frac{\{n+1\}}{\{n\}}$$

#### B cell clone lineage tree construction

Multi-compartmental clones were identified as B cell clones containing at least 1 sequence present in each of the paired peripheral blood and synovial samples from an individual donor. Lineage containing sequences that displayed evidence of index misassignment were discarded. Lineage trees of multi-compartmental clones were inferred using PHYLIP v3.6 [27] in Alakazam and graph layout was plotted in Gephi v0.9.2 using the ForceAtlas 2 algorithm [28].
